# Self-Incompatibility Alleles in Iranian Pear Cultivars

**DOI:** 10.1101/792507

**Authors:** Maryam Bagheri, Ahmad Ershadi

**Affiliations:** Department of Horticultural Sciences, Faculty of Agriculture, University of Bu-Ali Sina, Hamedan, Iran

**Keywords:** *Gametophytic self-incompatibility*, *Pyrus communis*, *PCR*, *S-alleles*, *Pear*

## Abstract

In the present study, the S-alleles of eighteen pear cultivars, (including fourteen cultivars planted commercially in Iran and four controls) are determined. 34 out of 36 S-alleles are detected using nine allele-specific primers, which are designed for amplification of S_101_/S_102_, S_105_, S_106_, S_107_, S_108_, S_109_, S_111_, S_112_ and S_114_, as well as consensus primers, PycomC1F and PycomC5R. S_104_, S_101_ and S_105_ were the most common S-alleles observed, respectively, in eight, seven and six cultivars. In 16 cultivars, (‘Bartlett’ (S_101_S_102_), ‘Beurre Giffard’ (S_101_S_106_), ‘Comice’ (S_104_S_105_), ‘Doshes’ (S_104_S_107_), ‘Koshia’ (S_104_S_108_), ‘Paskolmar’ (S_101_S_105_), ‘Felestini’ (S_101_S_107_), ‘Domkaj’ (S_104_S_120_), ‘Ghousi’ (S_104_S_107_), ‘Kaftar Bache’ (S_104_S_120_), ‘Konjoni’ (S_104_S_108_), ‘Laleh’ (S_105_S_108_), ‘Natanzi’ (S_104_S_105_), ‘Sebri’ (S_101_S_104_), ‘Se Fasleh’ (S_101_S_105_) and ‘Louise Bonne’ (S_101_S_108_)) both alleles are identified but in two cultivars, (‘Pighambari’ (S_105_) and ‘Shah Miveh Esfahan’ (S_107_)) only one allele is recognized. It is concluded that allele-specific PCR amplification can be considered as an efficient and rapid method to identify S-genotype of Iranian pear cultivars.

## Introduction

Self-incompatibility is a widespread genetic mechanism that inhibits self-fertilization in plants thus encourage out-crossing which has been frequently evolved among plants (de Nettancourt 1997). The top studied mechanisms of self-incompatibility take action by preventing the germination of pollen on stigmas, or the infiltration of the pollen tube in the styles. These mechanisms are based on protein-protein connections and are adjusted by a single locus named S, which has a lot of different alleles (Charlesworth et al. 2005). Self-incompatibility (SI) systems can be categorized into two main groups: gametophytic self-incompatibility (GSI) and sporophytic self-incompatibility (SSI). In GSI, the SI phenotype of the pollen is defined by its own gametophytic haploid genotype. If the male and female parents have totally different S-genotypes, i.e. they share no S-allele (e.g., S_1_S_2_ × S_3_S_4_), the cross is entirely compatible. If parents share one S-allele (e.g., S_1_S_2_ × S_1_S_3_) then pollen with the same allele is rejected and the cross is semi-compatible. Therefore, if parents have the same S-alleles at the self-incompatibility locus (e.g., S_1_S_2_ × S_1_S_2_) then pollen cannot grow on stigma. GSI system is the most common type of SI, existing in *Solanaceae*, *Rosaceae*, and *Papaveraceae* families (De Nettancourt 2001).

In SSI, the SI phenotype of the pollen is appointed by the diploid genotype of the anther. If the male and the female parent have one or two identical S-allele, (e.g., S_1_S_2_ × S_1_S_3_ or S_1_S_2_ × S_1_S_2_), then the growth of the pollen tube is prevented, and the cross is incompatible. Hence a successful cross will be possible if both parents do not have any equal S-allele (e.g., S_1_S_2_ × S_3_S_4_). This form of SI is found in *Brassicaceae*, *Asteraceae*, *Convolvulaceae*, *Betulaceae*, *Caryophyllaceae*, *Sterculiaceae* and *Polemoniaceae* families (De Nettancourt 2001).

Iranian pears belong to European pears (*Pyrus communis* L.), like other fruit species of the *Rosaceae* family, and exhibit gametophytic self-incompatibility system. In the *Rosaceae* S-lucas, a ribonuclease (S-RNase) is encoded and causes more progress in biochemical and molecular strategies for S-genotyping (Sanzol and Robbins 2008). Using alignment of the amino acid sequences of S-RNases, five conserved regions (C_1_, C_2_, C_3_, RC_4_, and C_5_), and one hypervariable (HV) region are specified. Also, a single intron, which is highly polymorphic, located between C_2_ and C_3_, within the HV is shown (Ushijima et al. 1998; De Franceschi et al. 2012). SFBB (S-locus F-box genes brothers) is another factor in GSI system controlling and is expressed in the pollen (De Franceschi et al. 2012).

Commercial fruit set in pear orchards appertains on cross-pollination and requires at least two compatible cultivars that flower simultaneously to permit fruit production (Zisovich et al. 2010). Consequently, S-genotype determination is essential for selection of compatible pollinizers and compatible crosses in breeding programs (Okada et al. 2015). Wherever scant numbers of parental lines are used in fruit breeding programs, narrow genetic base in new marketable cultivars is observed (Egea and Burgos 1996). For example, in the majority of breeding programs in European pear in the last decades, ‘Williams’ and ‘Coscia’ cultivars have been used as parental lines. So, repeated use of these cultivars leads to increment of cross-incompatibility cases in pears (Sanzol and Herrero 2002).

The traditional way of designation of compatibility between cultivars is time-consuming and expensive. Also, often it is affected by many other factors such as environmental and physiological parameters. It should be noted that it is complex to distinguish between fully compatible and semi compatible crosses (Zuccherelli et al. 2002). Recently, using of molecular methods for determination of S-genotype, where many of them are based on polymorphisms of the S-RNase gene, is increased (Sanzol and Robbins 2008). One exact way to determine S-genotype of pear cultivars is by identification of self-incompatibility alleles using allele-specific PCR amplification. This method is already used for identification of S-genotypes in almond (Ortega et al. 2006; Valizadeh and Ershadi 2009; Hafizi et al. 2013; Mousavi et al. 2014), apple (Broothaerts et al. 1995; Janssens et al. 1995; Ershadi et al. 2006), apricot (Vilanova et al. 2006; Murathan et al. 2017), sweet cherry (Szikriszt et al. 2013), Japanese pear (Ishimizu et al. 1999; Takasaki et al. 2004; Kim et al. 2007) and European pear (Moriya et al. 2007; Mota et al. 2007).

The first available report on the S-genotypes of European pear cultivars is published by Tomimoto et al. (1996), where they studied stylar proteins of European and Chinese pears and proposed various types of S-RNase for some pear cultivars. Using in vivo method, Sanzol and Herrero (2002) investigated the operation of pollen tube in pistil as well as pollen-pistil incompatibility in pears. However, it should be mentioned that these results are not in agreement with the previous allele designation used by Tomimoto et al. (1996).

Genomic PCRs with consensus primers, designed from nucleotide sequences of apple S-RNases, were used to amplify six putative S-RNase alleles in pears (Zuccherelli et al. 2002). Zisovich et al. (2004) identified seven S-alleles in European pear cultivars by PCR assay and sequencing, also expressed compatibility relations between nine pear cultivars.

Cleaved amplified polymorphic sequence (CAPS marker) system was established using primers annealing to S-locus sequences for genotyping European pear cultivars by Takasaki et al. (2006). In CAPS marker system, the resulting PCR fragments were digested with different restriction endonucleases (Takasaki et al. 2004; Mohring et al. 2005). In that study, full-length cDNAs of nine S-RNases from stylar RNA of five European pear cultivars were cloned (Takasaki et al. 2006). Using CAPS marker method, Moriya et al. (2007) found 17 S-alleles in 32 European pear cultivars, which were classified into 23 S-genotypes.

Mota et al. (2007) designed a consensus primer pair (Sall-F2 / Sall-R3) and determined S-genotypes of eight European pear cultivars (They used letter nomenclature). Sanzol et al. (2006) used number-based nomenclature instead of symbol-letter nomenclature for S-alleles appellation. Based on apple and Japanese pear conserved regions of S-RNase, Sanzol et al. (2006) designed degenerated S-RNase consensus primers (MPyC1F and MPyC5R) for *Pyrus communis*. These primers were amplified PCR products associated with alleles S_1_, S_3_, S_4_, and S_5_. Also, they designed a reverse primer (PycomS2R) for specific amplification of the S_1_ and S_2_ used in combination with MPyC5R. Later, Sanzol and Robbins (2008) made some changes in consensus primers (MPyC1F and MPyC5R) to amplify S_6_, S_7_, S_8_, S_9_, S_11_, and S_14_ S-RNases, in addition to the pervious alleles, but these primers could not amplify S_10_, S_12_, and S_13_ S-RNases. Therefore, they designed several new reverse primers to amplify S_10_, S_12_, and S_13_ alleles, in combination with MPyC1F. They, also, reported designing new consensus primers (PycomC1F and PycomC5R) using genomic sequences from C_1_ to C_5_ conserved regions that made them able to amplify all 14 S-RNase alleles. Moreover, they designed reverse allele specific primers for the S_5_, S_6_, S_7_, S_8_, S_9_, S_11_, S_12_, and S_14_ RNases using variable regions which could be used in combination with the PycomC1F consensus primer. Also, alternative forward primers were designed for S_7_ and S_8_. Sanzol and Robbins (2008) described that S-alleles genotypes resulted from PCR coincides exactly with the S-phenotypes inferred from the test of crosses. With increasing the number of studies on European pear S-genotypes, new S-alleles such as S_m_/S_n_/S_o_/S_q_/S_r_/S_20_/S_21_/S_23_ and S_24_ were identified and sequenced using PycomC1F and PycomC5R primers; Also, specific primers were designed for some of these new alleles (Sanzol 2009). Then, it was decided by Goldway et al. (2009) to renumber the European pear S-RNase alleles in order to distinguish between European and Japanese pear S-alleles. The new numeration started from 101 and subsequently, in the current study, the authors use this new system of the allele names.

No significant study heretofore has been carried out on Iranian pear cultivars. The aim of this paper is to determine S-genotypes of 14 Iranian and four European pear cultivars which are well adapted to Iranian climate. This study is performed via a molecular PCR based method (S-PCR) optimized with primers specifically designed for European pears. We were able to detect 34 out of 36 S-alleles of these cultivars.

## Materials and methods

### Plant materials and DNA extraction

14 important Iranian pear cultivars (‘Doshes’, ‘Koshia’, ‘Paskolmar’, ‘Flestini’, ‘Domkaj’, ‘Ghousi’, ‘Kaftar bache’, ‘Konjoni’, ‘Laleh’, ‘Natanzi’, ‘Pighambari’, ‘Sebri’, ‘Se Fasleh’ and ‘ShahMiveh’) and four European ones (‘Bartlett’, ‘Beurre Giffard’, ‘Comice’ and ‘Louise Bonne’) are used for identifying S-genotypes. The authors in this paper did not have access to a set of cultivars encompassing all known alleles. The available cultivars used as reference, i.e., ‘Bartlett’(S_101_S_102_), ‘Beurre Giffard’(S_101_S_106_), ‘Comice’(S_104_S_105_) and ‘Louise Bonne’ (S_101_S_102_) were previously genotyped by Sanzol et al. (2006) and Sanzol and Robbins (2008). Trees were located at Najaf Abaad and Mobarakeh fruit collection (Isfahan, Iran). During July, five to six leaves of young shoots of various cultivars were collected and immediately frozen and stored in liquid nitrogen, until the launch of DNA extraction. The extraction of DNA from leaves was based on the method described by Doyle and Doyle (1987) (CTAB method) with some modifications.

### S-RNase PCR amplification analysis

DNA samples were diluted to a final concentration of 10 ng.µL^−1^. PCR was performed using 50 ng of genomic DNA in a 30-µL reaction volume containing 1X reaction buffer (supplied with the enzyme), 2 mM MgCl_2_, 0.2 mM dNTPs, 0.6 mM of each primer and 0.8 U of Taq DNA polymerase. PCR amplification was conducted in an iCycler thermal cycler (Bio-Rad, Hercules, CA) with the following program: 2 min of denaturation at 94 ºC; 36 cycles of 30 s at 94 ºC, 1 min at specific annealing temperature, and 2 min at 72 ºC; and a final extension of 10 min at 72 ºC.

PCR specific S-allele amplifications in the pear cultivars were performed using two consensus primers PycomC1F (5′ ATTTTC AATTTACGCAGCAATATCAGC 3′) and PycomC5R (5′ CTGCAAAGWSHGACCTCAACCAATTC 3′) already designed and used for S-allele characterization in European pears by Sanzol and Robbins (2008) and later updated by Sanzol (2009). Using these consensus primers, alleles are amplified in the following sizes: S_101_ in 1300 bp, S_103_/S_112_ in 1600 bp, S_102_ in 1700 bp, S_4_ in 750 bp, S_110_ in 2200 bp, S_113_ in 2000 bp, S_106_/S_108_/S_111_/S_122_ in 675 bp, S_105_/S_107_/S_109_/S_114_/S_121_/S_123_/S_124_ in 650 bp and S_120_ in 800 bp. The specific primers designed by Sanzol and Robbins (2008) were used to amplify the S_101/102_, S_105_, S_106_, S_107_, S_108_, S_109_, S_111_, S_112_ and S_114_ alleles (Table 1).

**Table 1.**
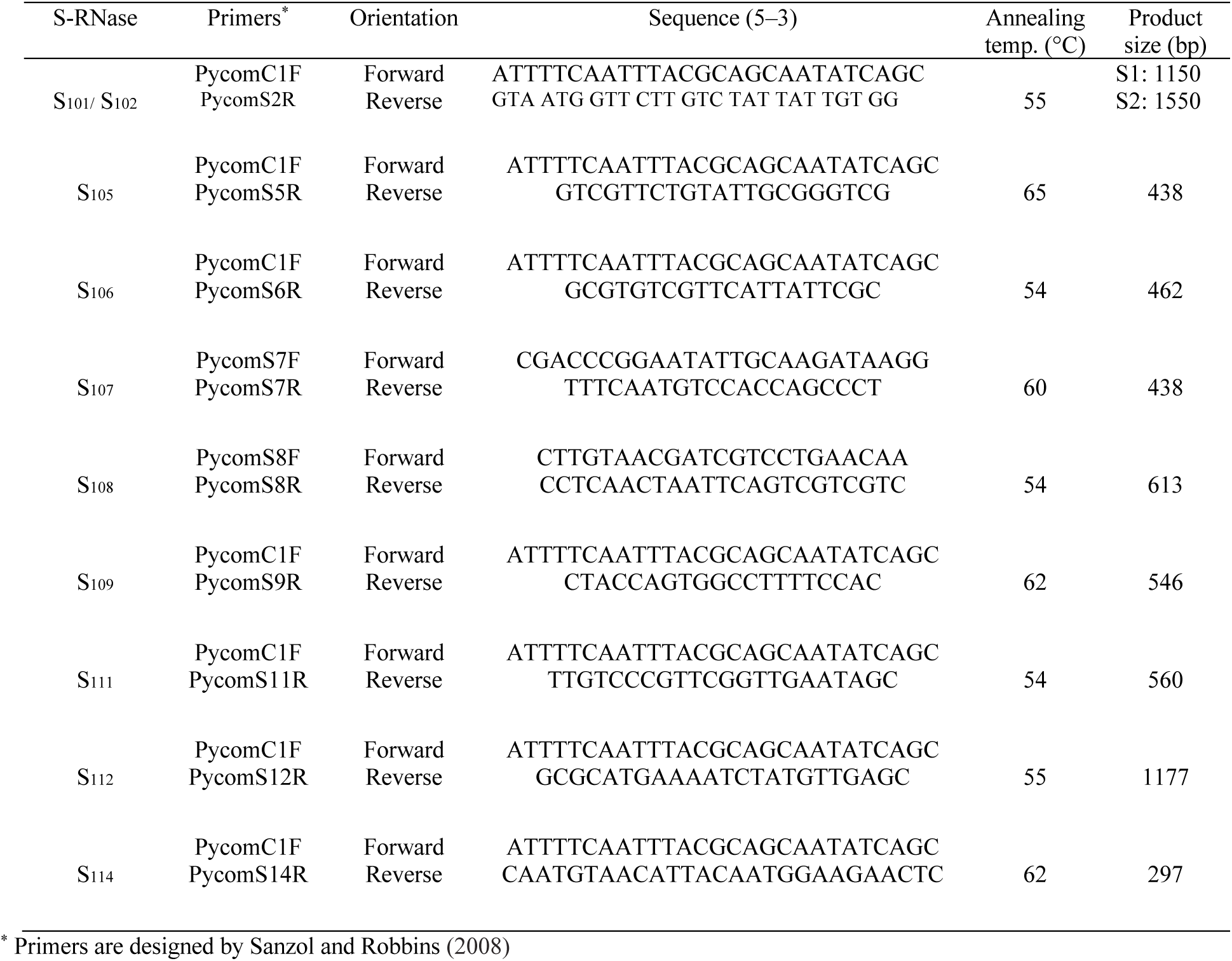
Primers and reaction conditions for allele-specific PCR of S-RNases in European pear.

Annealing temperature for each pair of primers were determined using gradient. All PCR reactions were executed at least two times. The molecular sizes of the PCR products were estimated using 100 bp Plus DNA ladder. The amplified products were separated in 1.5% (w/v) agarose gels in 1X TAE buffer at 60 V for 4 h. After electrophoresis, the gels were stained with ethidium bromide and visualized under the UV light using UVitec gel documentation.

## Results and discussion

### Confirmation with allele-specific primers

Using the mentioned nine specific primer pairs (Table 1), six self-incompatibility alleles, including S_101_, S_102_ (by the same primer), S_105_, S_106_, S_107_ and S_108_, were amplified in the studied cultivars and four alleles, including S_109_, S_111_, S_112_ and S_114_, were not amplified in none of these studied cultivars. Also, S_102_ and S_106_ could not be found among Iranian cultivars. Two alleles were determined in seven cultivars and only one allele was determined in nine cultivars. No allele was found in the remaining two cultivars, namely ‘Domkaj’ and ‘Natanzi’, using these primers (Fig. 1 and Table 2).

**Table 2.**
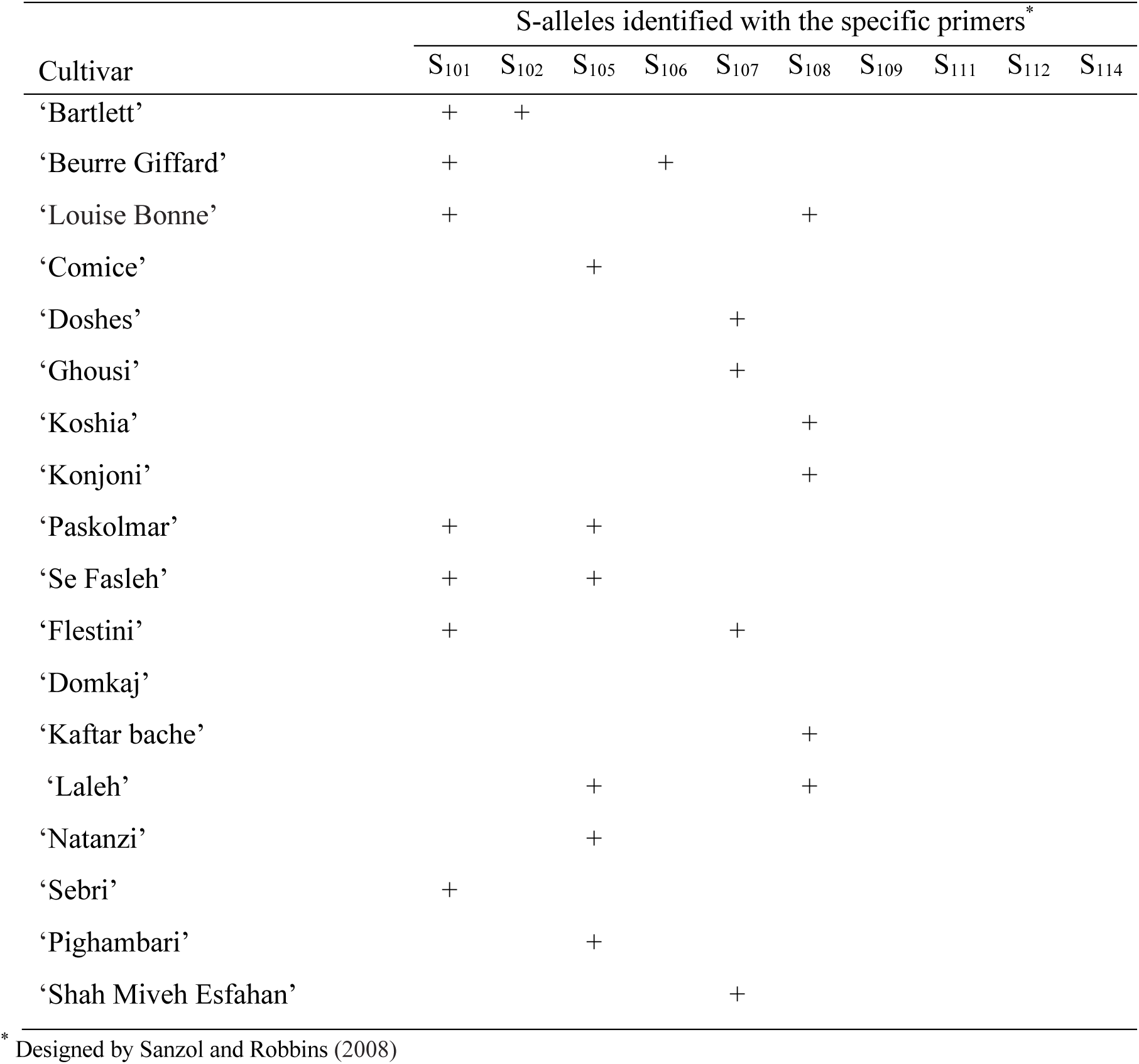
S-alleles identified in 14 pear cultivars plus four controls using allele-specific primers.

**Fig. 1.**
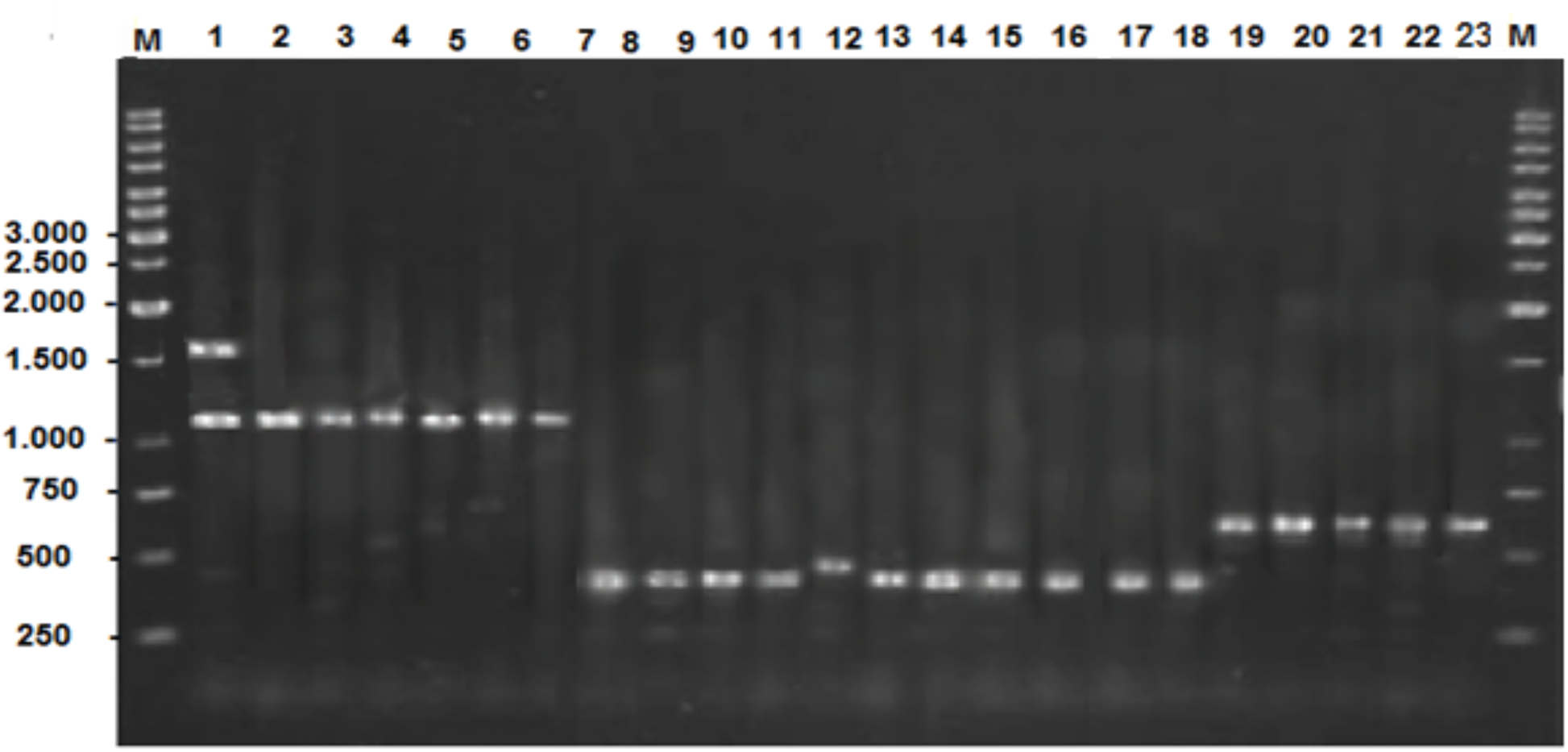
PCR amplification of S-alleles of 14 Iranian and four reference pear cultivars using allele-specific primers. Lanes, M: 100 bp Plus DNA ladder; 1 – 23: S_101/102_: ‘Bartlett’, ‘Beurre Giffard’, ‘Louise Bonne’, ‘Paskolmar’, ‘Se Fasleh’, ‘Flestini’, ‘Sebri’. S_107_: Doshes’, ‘Ghousi’, ‘Flestini’, ‘Shah Miveh’. S_106_: ‘Beurre Giffard’. S_105_: ‘Comice’, ‘Paskolmar’, ‘Se Fasleh’, ‘Laleh’, ‘Pighambari’, ‘Natanzi’. S_108_: ‘Louise Bonne’, ‘Koshia’, ‘Konjoni’, ‘Kaftar bache’, ‘Laleh’. S_120_: ‘Domkaj’ and ‘Kaftar bache’; M: 100 bp Plus DNA ladder.

S-genotypes were recognised in three European pear cultivars, including ‘Bartlett’ (S_101_S_102_), ‘Beurre Giffard’ (S_101_S_106_) and ‘Louise Bonne’ (S_101_S_108_), using allele-specific polymerase chain reaction. Also, S-genotypes were detected in four Iranian cultivars, ‘Flestini’ (S_101_S_107_), ‘Se Fasleh’ (S_101_S_105_), ‘Paskolmar’ (S_101_S_105_), and ‘Laleh’ (S_105_S_108_) (Fig. 1 and Table 2).

After identification of S-genotypes of pear cultivars by specific primers, it was seen that the present results about ‘Bartlett’ (S_101_S_102_) cultivar were equal to the results given by Sanzol and Robbins (2008), Takasaki et al. (2006) and Mota et al. (2007). Moreover, present findings on S-genotype in ‘Beurre Giffard’ (S_101_S_106_) cultivar were similar to those given by Moriya et al. (2007) and Sanzol and Robbins (2008). Detected S-genotype for ‘Louise Bonne’ (S_101_S_108_) cultivar, by the current study, was different from the one shown by Sanzol and Robbins (2008), which may be due to different cultivars with the same name in different regions.

### Confirmation with consensus primers

Sanzol and Robbins (2008) used partial genomic sequences of the S-RNase gene, spanning between the C_1_ and C_5_ conserved regions. They designed a pair of consensus primers (PycomC1F and PycomC5R) for 14 S-alleles (named S_101_-S_114_). Also, Sanzol (2009) detected that other five S-alleles (named S_120_-S_124_) can be amplified with these consensus primers. Using these primers, we identified both S-alleles for 16 studied pear cultivars (12 Iranian cultivars and four references - European - ones), but only one allele was detected in the other two Iranian cultivars, i.e., ‘ShahMiveh’ and ‘Pighambari’, where only one band with size 650 bp was amplified (Table 3).

**Table 3.** Sizes of amplification products and S-alleles identified in 18 Iranian and European pear cultivars using consensus primers (PycomC1F and PycomC5R designed by Sanzol and Robbins, 2008).

| Cultivar | Sizes of genomic PCR products (bp) |
| --- | --- |
| ‘Bartlett’ | 1300,1700 |
| ‘Beurre Giffard | 1300,675 |
| ‘Louise Bonne’ | 1300,675 |
| ‘Comice’ | 650,750 |
| ‘Doshes’ | 750,650 |
| ‘Ghousi’ | 750,650 |
| ‘Koshia’ | 750,675 |
| ‘Konjoni’ | 750,675 |
| ‘Paskolmar’ | 1300,650 |
| ‘Se Fasleh’ | 1300,650 |
| Flestini’ | 1300,650 |
| ‘Domkaj’ | 750,800 |
| ‘Kaftar bache’ | 675,800 |
| ‘Laleh’ | 650,675 |
| ‘Natanzi’ | 650,750 |
| ‘Sebri’ | 1300,750 |
| ‘Pighambari’ | 650 |
| ‘Shah Miveh Esfahan’ | 650 |

Figure 2 shows the amplification of self-incompatibility alleles in the studied pear cultivars, using consensus primers designed by Sanzol and Robbins (2008) on 1.5% agarose gel. All data obtained using specific primers were completely verified using consensus primers (Tables 3 and 4).

**Table 4.**
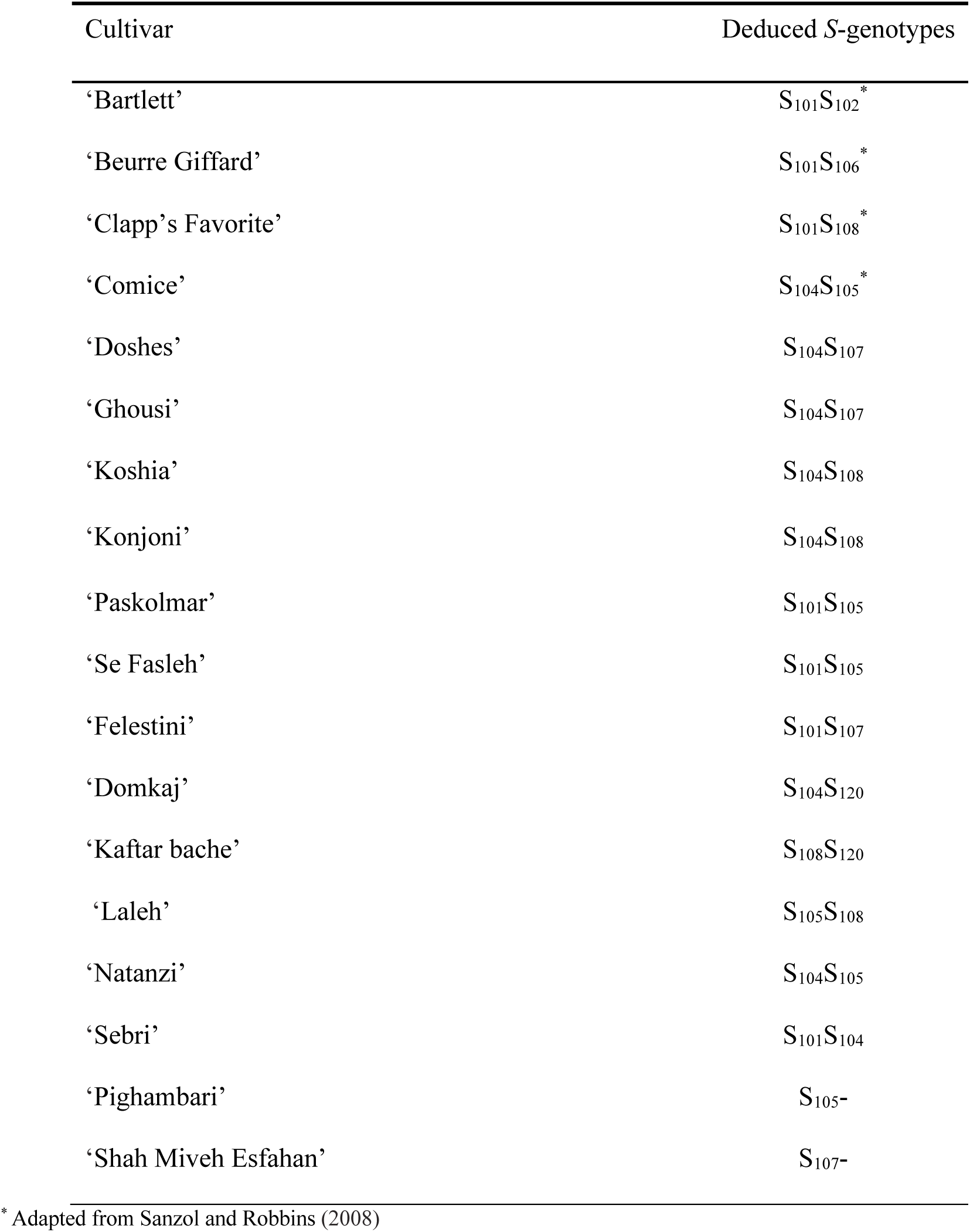
S-genotypes deduced in 18 pear cultivars obtained with consensus and allele-specific primers.

**Fig. 2.**
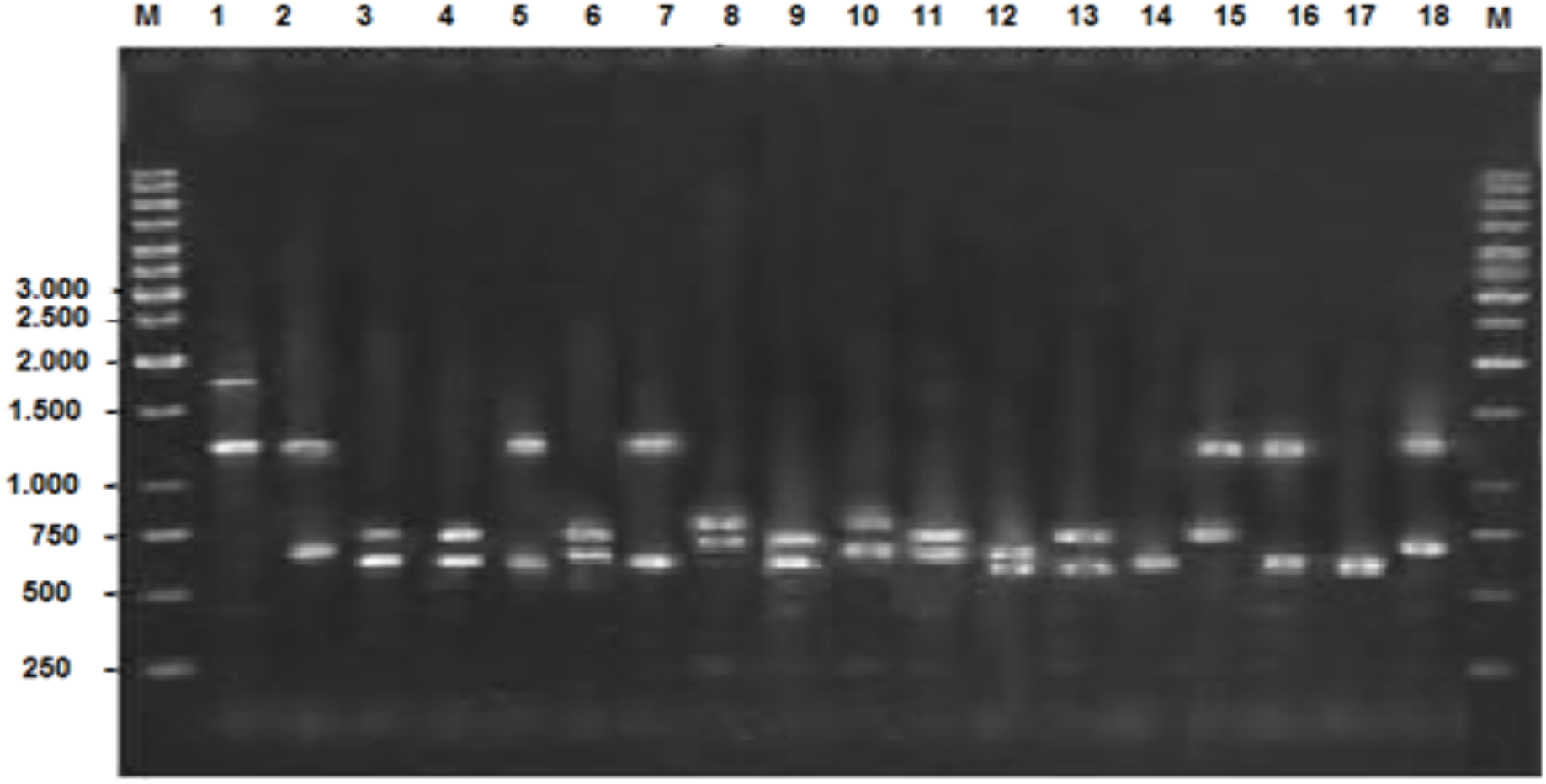
PCR amplification of *S*-alleles in DNA from 14 Iranian and 4 none Iranian pear cultivars using consensus primers. Lanes M: 100 bp Plus DNA ladder; 1 – 18: Bartlett’, Beurre Giffard’, ‘Comice’, ‘Doshes’, ‘Flestini’, ‘Koshia’, ‘Paskolmar’, ‘Domkaj’, ‘Ghousi’, ‘Kaftar bache’, ‘Konjoni’, ‘Laleh’, ‘Natanzi’, ‘Pighambari’, ‘Sebri’, ‘Se Fasleh’, ‘Shah Miveh’, ‘Louise Bonne’; M: 100 bp Plus DNA ladder.

The results obtained using specific primers indicated that the amplified band, is S_105_, in ‘Pighambari’ cultivar, and is S_107_, in ‘ShahMiveh’ cultivar. However, we could not explain their S-genotype. It is possible that both alleles were amplified with similar size; Or, the non-detected band could be produced with a large size that is not recognized. In this case, the identification of the new S-allele, needs another, or a modified, primer.

## Conclusion

In this study, the S-genotypes of 18 pear cultivars (planted in Iran) were determined with different sets of consensus and allele-specific primers (Table 4). Using these primers, the S-genotypes of 16 cultivars were identified. S_104_, S_101_, and S_105_ were the most common alleles found, respectively, in eight, seven and six cultivars assayed in this study.

The ‘Kaftar bache’ (S_108_S_120_), ‘Domkaj’ (S_104_S_120_) and ‘Laleh’ (S_105_S_108_) cultivars-can be categorized in common pollinizers, as they have a special incompatibility genotype and, so far, similar incompatibility genotype has not been reported for them.

Investigation on the self-incompatibility alleles can be a good indicator of the differences between Iranian pears and European cultivars. The incompatibility genotypes have been detected by consensus and specific primers. However, all self-incompatibility alleles were not identified by using the method of allele-specific amplification by PCR, but this method is an applicable tool for young seedlings. Moreover, it is a fast and cheap method, meaning that with a small amount of leaf or any part of the plant, incompatibility genotypes can be investigated. It also carefully examines the self-incompatibility of fruit trees. However, molecular genotyping has several intrinsic limitations that should not be ignored. First, it may be insensitive toward unknown S-alleles; Therefore, it is important to complement these methods with a reliable and efficient in-vivo test to control putative new specificities.

This information assists nurseries, growers, and breeders to select compatible combinations of cultivars. According to our results, allele-specific PCR amplification is an efficient and rapid method to identify S-genotype of Iranian pear cultivars. The use of different sets of consensus and allele-specific primers results in the complete and unequivocal identification of S-genotypes in most commercial Iranian pear cultivars. In the case of designing the specific primers for other self-incompatibility alleles, S-genotype can be identified with higher speed and accuracy.

## Data Archiving Statement

In this study, there are no new data for submission to a database. Accession numbers at the NCBI database related to all the mentioned S-alleles are available in file S1.

## References

Broothaerts W, Janssens GA, Proost P, Broekaert WF (1995) cDNA cloning and molecular analysis of two self-incompatibility alleles from apple. Plant molecular biology 27(3):499–511.

Charlesworth D, Vekemans X, Castric V, Glémin S (2005) Plant self-incompatibility systems: a molecular evolutionary perspective. New Phytologist 168(1):61–69.

De Franceschi P, Dondini L, Sanzol J (2012) Molecular bases and evolutionary dynamics of self-incompatibility in the Pyrinae (Rosaceae). Journal of experimental botany 63(11):4015–4032.

De Nettancourt D (1997) Incompatibility in angiosperms. Sexual Plant Reproduction 10(4):185–199.

De Nettancourt D (2001) Incompatibility and incongruity in wild and cultivated plants (Vol. 3). Springer Science & Business Media.

Doyle JJ, Doyle JL (1987) A rapid DNA isolation procedure for small quantities of fresh leaf tissue. Phytochemical Bulletin 19:11–15.

Egea J, Burgos L (1996) Detecting cross-incompatibility of three North American apricot cultivars and establishing the first incompatibility group in apricot. Journal of the American Society for Horticultural Science 121(6):1002–1005.

Ershadi A, Talaii A (2006) Identification of S-alleles in 40 apple (Malus x domestica BORKH) cultivars by allele-specific PCR amplification. In XXVII International Horticultural Congress-IHC 2006: II International Symposium on Plant Genetic Resources of Horticultural 760:111–116.

Goldway M, Takasaki-Yasuda T, Sanzol J, Mota M, Zisovich A, Stern RA, Sansavini S (2009) Renumbering the S-RNase alleles of European pears (Pyrus communis L.) and cloning the S109 RNase allele. Scientia Horticulturae 119(4):417–422.

Hafizi A, Shiran B, Maleki B, Imani A, Banović B (2013) Identification of new S-RNase self-incompatibility alleles and characterization of natural mutations in Iranian almond cultivars. Trees 27(3):497–510.

Ishimizu T, Inoue K, Shimonaka M, Saito T, Terai O, Norioka S (1999) PCR-based method for identifying the S-genotypes of Japanese pear cultivars. Theoretical and Applied Genetics 98(6-7):961–967.

Janssens GA, Goderis IJ, Broekaert WF, Broothaerts W (1995) A molecular method for S-allele identification in apple based on allele-specific PCR. Theoretical and Applied Genetics 91(4):691–698.

Kim H, Kakui H, Koba T, Hirata Y, Sassa H (2007) Cloning of a new S-RNase and development of a PCR-RFLP system for the determination of the S-genotypes of Japanese pear. Breeding Science 57(2):159–164.

Mohring S, Horstmann V, Esch E (2005) Development of a molecular CAPS marker for the self-incompatibility locus in Brassica napus and identification of different S alleles. Plant breeding 124(2):105–110.

Moriya Y, Yamamoto K, Okada K, Iwanami H, Bessho H, Nakanishi T, Takasaki T (2007) Development of a CAPS marker system for genotyping European pear cultivars harboring 17 S alleles. Plant cell reports 26(3):345–354.

Mota M, Tavares L, Oliveira CM (2007) Identification of S-alleles in pear (Pyrus communis L.) cv. ‘Rocha’and other European cultivars. Scientia horticulturae 113(1):13–19.

Mousavi A, Babadaei R, Fatahi R, Zamani Z, Dicenta F, Ortega E (2014) Self-incompatibility in the Iranian Almond Cultivar ‘Mamaei’Using Pollen Tube Growth, Fruit Set and PCR Technique.

Murathan ZT, Kafkas S, Asma BM (2017) Determination of S alleles in Paviot x Levent apricot progenies by PCR and controlled pollination. Journal of applied botany and food quality 90:147–153.

Okada N, Sanada Y, Hirata Y, Yamada N, Wakiya T, Ihara Y, et al. (2015) The impact of rituximab in ABO-incompatible pediatric living donor liver transplantation: The experience of a single center. Pediatric transplantation 19(3):279–286.

Ortega E, Bošković RI, Sargent DJ, Tobutt KR (2006) Analysis of S-RNase alleles of almond (Prunus dulcis): characterization of new sequences, resolution of synonyms and evidence of intragenic recombination. Molecular Genetics and Genomics 276(5):413–426.

Sanzol J (2009) Genomic characterization of self-incompatibility ribonucleases (S-RNases) in European pear cultivars and development of PCR detection for 20 alleles. Tree genetics & genomes 5(3):393–405.

Sanzol J, Herrero M (2002) Identification of self-incompatibility alleles in pear cultivars (Pyrus communis L.). Euphytica 128(3):325–331.

Sanzol J, Robbins TP (2008) Combined analysis of S-alleles in European pear by pollinations and PCR-based S-genotyping; correlation between S-phenotypes and S-RNase genotypes. Journal of the American Society for Horticultural Science 133(2):213–224.

Sanzol J, Sutherland BG, Robbins TP (2006) Identification and characterization of genomic DNA sequences of the S-ribonuclease gene associated with self-incompatibility alleles S1 to S5 in European pear. Plant Breeding 125(5):513–518.

Szikriszt B, Doğan A, Ercisli S, Akcay ME, Hegedűs A, Halász J (2013) Molecular typing of the self-incompatibility locus of Turkish sweet cherry genotypes reflects phylogenetic relationships among cherries and other Prunus species. Tree genetics & genomes 9(1):155–165.

Takasaki T, Moriya Y, Okada K, Yamamoto K, Iwanami H, Bessho H, Nakanishi T (2006) cDNA cloning of nine S alleles and establishment of a PCR-RFLP system for genotyping European pear cultivars. Theoretical and Applied Genetics 112(8):15–43.

Takasaki T, Okada K, Castillo C, Moriya Y, Saito T, Sawamura Y, et al. (2004) Sequence of the S9-RNase cDNA and PCR-RFLP system for discriminating S1-to S9-allele in Japanese pear. Euphytica 135(2):157.

Tomimoto Y, Nakazaki T, Ikehashi H, Ueno H, Hayashi R (1996) Analysis of self-incompatibility-related ribonucleases (S-RNases) in two species of pears, Pyrus communis and Pyrus ussuriensis. Scientia Horticulturae 66(3-4):159–167.

Ushijima K, Sassa H, Tao R, Yamane H, Dandekar AM, Gradziel TM, Hirano H (1998) Cloning and characterization of cDNAs encoding S-RNases from almond (Prunus dulcis): primary structural features and sequence diversity of the S-RNases in Rosaceae. Molecular and General Genetics MGG 260(2-3):261–268.

Valizadeh B, Ershadi A (2009) Identification of self-incompatibility alleles in Iranian almond cultivars by PCR using consensus and allele-specific primers. The Journal of Horticultural Science and Biotechnology 84(3):285–290.

Vilanova S, Badenes ML, Burgos L, Martínez-Calvo J, Llácer G, Romero C (2006) Self-compatibility of two apricot selections is associated with two pollen-part mutations of different nature. Plant physiology 142(2):629–641.

Zisovich AH, Raz A, Stern RA, Goldway M (2010) Syrian pear (Pyrus syriaca) as a pollinator for European pear (Pyrus communis) cultivars. Scientia horticulturae, 125(3):256–262.

Zisovich A, Stern R, Shafir S, Goldway M (2004) Identification of seven S-alleles from the European pear (Pyrus communis) and the determination of compatibility among cultivars. The Journal of Horticultural Science and Biotechnology 79(1):101–10.

Zuccherelli S, Broothaerts W, Tassinari P, Tartarini S, Dondini L, Bester A, Sansavini S (2002) S-allele characterization in self-incompatible pear (Pyrus communis): Biochemical, molecular and field analyses. Acta Horticulturae 1:147–152.

